# PANDORA: a fast, anchor-restrained modelling protocol for peptide:MHC complexes

**DOI:** 10.1101/2022.03.04.482467

**Authors:** Farzaneh Meimandi Parizi, Dario F. Marzella, Derek Van Tilborg, Nicolas Renaud, Daan Sybrandi, Rafaella Buzatu, Li C Xue

**Affiliations:** Center for Molecular and Biomolecular Informatics, Radboudumc, Nijmegen, The Netherlands; Institute of Biochemistry and Biophysics, University of Tehran, Tehran, Iran; Eindhoven University of Technology, Dept. Biomedical Engineering, Institute for Complex Molecular Systems, Eindhoven, The Netherlands; Netherlands eScience Center, Amsterdam, The Netherlands; Bijvoet Centre for Biomolecular Research, Faculty of Science - Chemistry, Utrecht University, Utrecht, The Netherlands

**Keywords:** peptide-MHC, Integrative modelling, Computational Structural Biology, computational immunology, large-scale modelling

## Abstract

Deeper understanding T-cell mediated adaptive immune responses bears important implications in designing cancer immunotherapies and antiviral vaccines against pandemic outbreaks. T cells fire when they recognize foreign peptides that are presented on the cell surface by Major Histocompatibility Complexes (MHC), forming peptide:MHC (pMHC) complexes. 3D structure of pMHC complexes provides fundamental insight of T-cell recognition mechanism and aids immunotherapy design. High MHC and peptide diversities necessitate efficient computational modelling to enable whole proteome structural analysis.

Here we present PANDORA, a robust pMHC structure modelling pipeline. Given a query, PANDORA searches for structural templates in its extensive database and then applies anchor restraints to the modelling process. This restrained energy minimization ensures one of the fastest pMHC modelling pipelines so far. On a set of 835 pMHC-I complexes over 78 MHC types, PANDORA generated models with a median RMSD of 0.68 Å and achieved a 93% success rate in top 10 models. PANDORA performs competitively with three pMHC-I modelling state-of-the-art approaches and outperforms AlphaFold in terms of accuracy while being superior to it in speed. PANDORA is a modularized and user-configurable python package with easy installation. We envision PANDORA to fuel deep learning with large-scale high-quality 3D models to tackle long-standing immunology challenges.

## 1 Introduction

The adaptive T cell immune activation requires the T-cell Receptor (TCR) to recognize foreign peptide antigens presented on self MHC (Major Histocompatibility Complex) molecules. Intracellular antigenic peptides presented by MHC-I can activate CD8+ T cells, which can directly kill infected cells that present these peptides on their surface. Exogenously derived peptide antigens presented by MHC-II can activate CD4+ T cells, which stimulate the production of antibodies and promote CD8+ T Cell response (1).

Structural investigations of pMHC have provided profound knowledge of MHC antigen-display mechanisms and T cell functions (2). Such knowledge can be used to aid the design of new therapies for cancer (3), viral infections (4,5), and autoimmune disorders (6,7). MHC is the most polymorphic protein known to date in humans. MHC has more than 10,000 identified alleles (8). Each of these alleles has a specific binding preference for different peptides. Regardless of the highly polymorphic nature of MHC sequence, the MHC structure has been elucidated as having an “Ultra-conserved” fold (8). Considering the structure of MHC-I molecules, the peptide binding groove is formed by an a-chain, which has two domains denoted as G-ALPHA1 and G-ALPHA2 in IMGT nomenclature (9)(Figure 1A). This cleft is closed on the sides and contains two deep binding pockets. Short peptides of around 8 to 11 residue lengths canonically bind to these pockets with their second (P2) and last (PΩ) residues, respectively (10,11). The peptide-binding groove of MHC-II is formed by two domains from a- and /)-chain (corresponding to the G-ALPHA and G-BETA domains in IMGT nomenclature). Its binding cleft is open in both ends, and hence it accommodates longer peptides (Figure 1B). Normally 9 residues of the peptide bind to MHC binding groove, called binding core and the rest of the peptide protrude out of the groove. The peptide is anchored at the P1, P4, P6 and P9 pockets of MHC-II (11).

**Figure 1.**
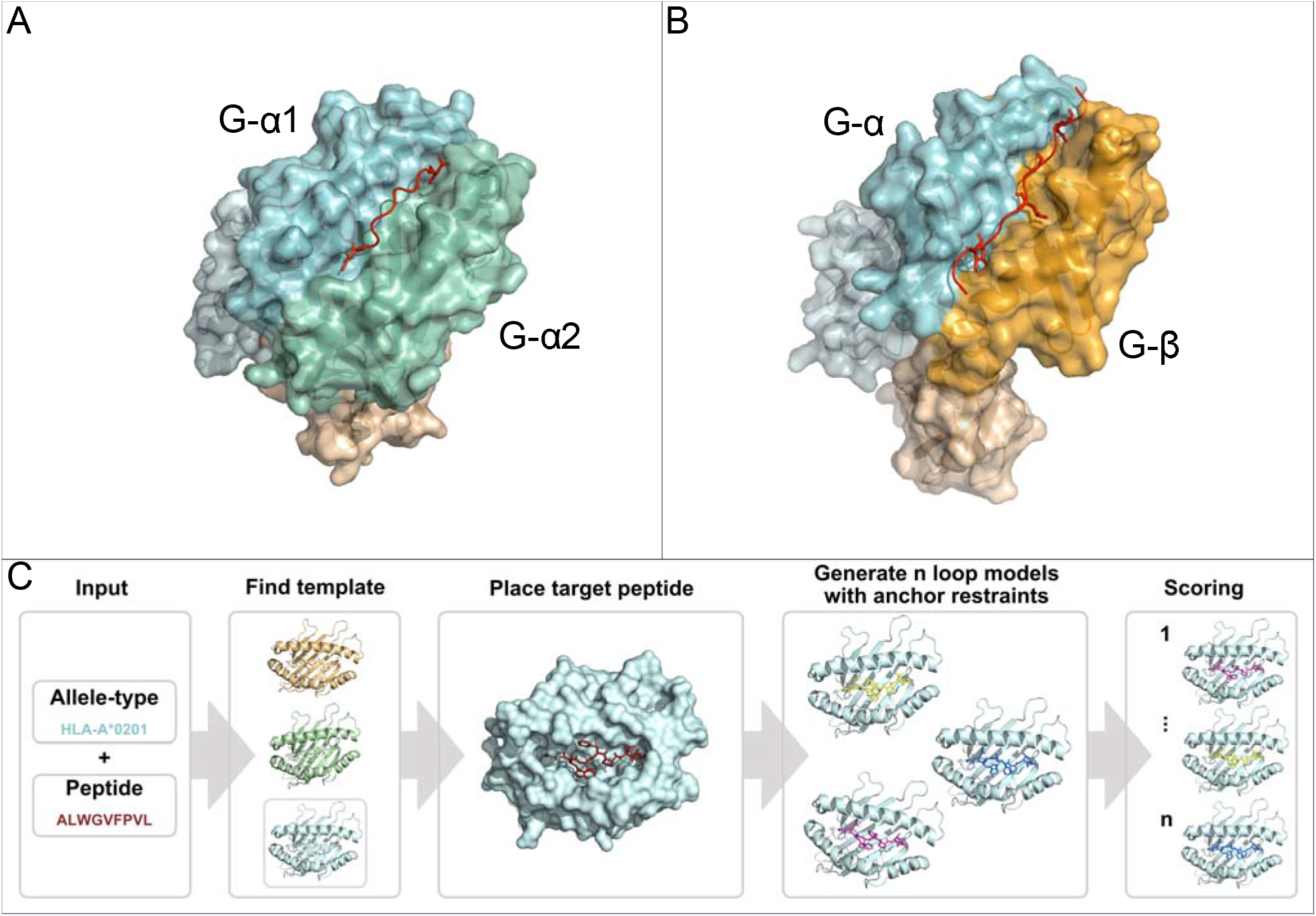
Overview of the MHC molecules and modelling process of PANDORA. **A)** 3D structure of a pMHC I complex (PDB ID: 1DUZ). The α chain is divided in IMGT defined domains by shades of light blue. The /J-2 Microglobulin chain is shown in light orange. The peptide is shown in red. **B)** 3D structure of a pMHC II complex (PDB ID: 1AQD). The alpha chain is divided in IMGT defined domains by shades of light blue. The /J chain is divided in IMGT defined domains by shades of orange. The peptide is shown in red. **C)** PANDORA schematic flowchart. An allele type and peptide sequence of a target pMHC-I case are given as input. This information is used to identify the best matching template structure from a local database of p:MHC-I structures. The target MHC is then modelled on top of the template and its peptide (red) is superposed on the template peptide. The anchor positions (specified by the user or by other tools, see section 2.4) are then specified as fixed. MODELLER then generates 20 loop models maintaining the anchor restrained. Finally, all models are scored with MODELLER internal molpdf scoring function.

Complementary to atomic-resolution 3D structure determination experiments (such as X-ray and NMR), the recent advances of large-scale mass spectrometry provide valuable tools to detect MHC binding peptides (12–14). However, an infinite number of potential peptides could be derived from host cells and diverse pathogens. The high diversity of MHC and peptide sequences call for the development of effective computational methods for predicting MHC-binding peptides and their binding affinities.

Currently, sequence-based machine learning (ML) methods are dominating MHC epitope predictions and outperform three-dimensional (3D) modelling methods (15–18). But their sequence-nature makes them incapable of incorporating key knowledge about 3D structures. Further, state-of-the-art ML approaches heavily rely on training data, and thus may have limited prediction power on rare MHC alleles. Alternatively, free energy calculation methods have shown several successful cases in estimating binding affinity changes upon mutations (19), but such methods are not computationally feasible for large-scale proteome scanning. Even so, 3D modelling approaches have several compelling theoretical advantages: (1) they are sensitive to the minor changes in the 3D space- and energy-landscape caused by mutations (20), (2) they exploits fundamental physics-based energies, thus could potentially be generalizable to many other MHC alleles with limited binding affinity data, and (3) they offer 3D models of MHC:peptide and TCR:peptide:MHC, which could be used to guide experimental work.

In the past decades, many efforts have been devoted to design reliable modelling approaches to model 3D structures of pMHC complexes. There are three basic approaches for modelling 3D pMHC structures: (1) molecular dynamics (MD) (21–23), (2) molecular docking (24–26), and (3) homology modelling (27) (see review (28)). MD approaches have shown to produce accurate structures; however, they are computationally intensive. State-of-the-art methods are often hybrid methods of these three techniques to make pMHC modelling computationally accessible while being reliable (27,29,30). The general design of the state-of-the-art methods is as: (i) generating peptide conformation(s) based on a template conformation, (ii) inserting peptide into fixed MHC-I backbone, and (iii) optimization of the overall conformation including side-chains.

Several state-of-the-art methods for modelling pMHC-I are available. DockTope (30) models pMHC-I complexes for 4 different MHC-I allotypes. It docks the peptide to MHC-I with different initial points and then selects the best conformation. It subsequently optimizes the conformation of pMHC-I with GROMACS (31) and repeats the docking to refine the pMHC-I structure. GradDock (29) as the other pipeline, constrains the peptide ends and generates numerous peptide conformations, next uses steered gradient descent to simulate binding of the peptide to MHC-I. After topological correction, a novel Rosetta-based scoring function selects the best candidate. Later APE-Gen (27) was proposed, adding the receptor modelling with MODELLER (32) before the main mentioned steps. APE-Gen also anchors the peptide termini and utilizes Random Coordinate Descent (RCD) (33) loop modelling to generate 100 peptide conformations. For energy optimization, it utilizes a molecular docking tool. In APE-Gen, all the main steps are run iteratively.

More recently, AlphaFold (34) and RoseTTAfold (35) have shown to have outstanding results in single-chain protein structure prediction. In this regard, there have been few attempts modelling peptides using AlphaFold, either using AlphaFold multimer (36) or linking the peptide to the protein using peptide linkers (37,38). However, peptide:MHC interactions present unique challenges and have not been solved yet. This is mainly due to two factors: 1) peptides in peptide:MHC databases often have synthetic or frameshift origin, meaning they can have not enough evolutionary information to generate an MSA (Multiple Sequence Alignment), which is the main piece of knowledge used as input from these DL-based prediction methods (34). 2) peptides are highly flexible; therefore, the use of specific domain knowledge is essential to reduce the large conformational space for speed (e.g. properly guiding the anchors docking/modelling of peptide:MHC). General purpose AI software is often slower than integrative modelling, thus not fitting to model millions of peptide:MHC interactions.

Here we present PANDORA (Peptide ANchoreD mOdelling fRAmework), a user-friendly, fast, and reliable pMHC integrative modelling pipeline. PANDORA leverages two key structural concepts of the interaction of MHC-I-peptide binding: first, MHC molecules having a highly conserved overall structure while the binding cleft surface is highly variable; second, MHCs use several anchor pockets (2 for MHC-I and 4 for MHC-II) to dock peptides. PANDORA first builds a homology model of the MHC structure by using a similar allele template selected for each target. Then, it aligns the target anchors with the template anchors and restraints them to reduce the conformational search space. Finally, PANDORA performs a loop energy minimization to produce an accurate model of the peptide conformation. By using a restrained energy minimization step, the modelling phase is kept short, resulting in one of the fastest pMHC modelling pipelines so far. This enables large-scale proteome modelling of pMHCs for training subsequent ML algorithms.

We first demonstrate PANDORA’s performances on a cross-validation set of 835 pMHC class I structures. We then compare it with three pMHC-I modelling softwares on several pMHC sets with experimental structures. Finally, we performed a qualitative evaluation of different AlphaFold approaches against PANDORA on modelling 4 pMHC-I structures. PANDORA performs competitively, or better than, these pMHC-I modelling softwares in terms of accuracy and computational time while providing an easier installation and flexible user experience. PANDORA is efficient enough for large-scale modelling of pMHC complexes.

## 2 Results

### 2.1 Description of PANDORA

Our information-driven homology modelling framework PANDORA takes a few crucial steps (Figure 1C) to provide core domain knowledge to MODELLER (32). MHC high structural homology and anchoring positions for bound peptides are used to constrain the conformational search space to effectively produce an ensemble of 3D models.

First, PANDORA builds a large template database, which consists of all valid peptide-MHC structures from IMGT/3Dstructure-DB. All structures in the template database are renumbered starting from 1. The renamed chain ID of peptides is P; that of MHC is M (M and N for MHC-II alpha and beta chain, respectively). As allele names, PANDORA relies on G-domain allele names from IMGT, which are assigned based on Multiple Sequence Alignments of only G-domains, because this is the domain responsible for the peptide binding (9). One structure can have more than one allele name since the same G-domain can be shared by multiple MHC alleles. For this reason, any reference to MHC allele further in this paper has to be intended to G-domain alleles.

During a modelling run, PANDORA selects a suitable structural template for the given target from our parsed database. It then uses MODELLER to build an initial 3D structure, keeps the anchors restrained and performs a loop modelling on the central region of the peptide to output the final structures. Output models are ranked to indicate to the user which are the best ones.

### 2.2 PANDORA produces near-native models on a large benchmark set

We benchmarked PANDORA on all pMHC-I complexes with experimentally determined structures in the IMGT/3Dstructure-DB database (39) (as of 28/06/2021): 835 complexes over 78 MHC allele types (PDB IDs reported in Suppl. Table 1). We removed one structure from this dataset and used it as the test case. This process was repeated for every structure in the dataset. We used MHC allele name, actual peptide anchor position and peptide sequence as inputs for PANDORA, and asked PANDORA to generate 20 model structures.

When evaluating on the lowest Ligand Root Mean Square Deviation (L-RMSD) (40) obtained per case, our pipeline is capable of producing at least one near-native model (L-RMSD < 2L) in 96.4% of the cases and an overall mean deviation of 0.81 ± 0.56 L (Figure 2A). MODELLER’s internal molpdf function provides high-quality ranking for the models produced by PANDORA. To select which models should be provided to the user as output, we evaluated MODELLER’s scoring functions molpdf and DOPE with compared hit rate and success rate plots (Suppl. Fig. 1), obtaining the best results from molpdf. Figure 2C shows this scoring function reaching a success rate of 93% in the top 10 models (see Methods section 5.3). The resulting L-RMSD median of top scored models with molpdf (the final best output provided to the user) is 0.85 Å. Therefore, PANDORA’s scoring method, together with the sampling procedure, allows us to deliver reliable predictions (Figure 2B and D).

**Figure 2.**
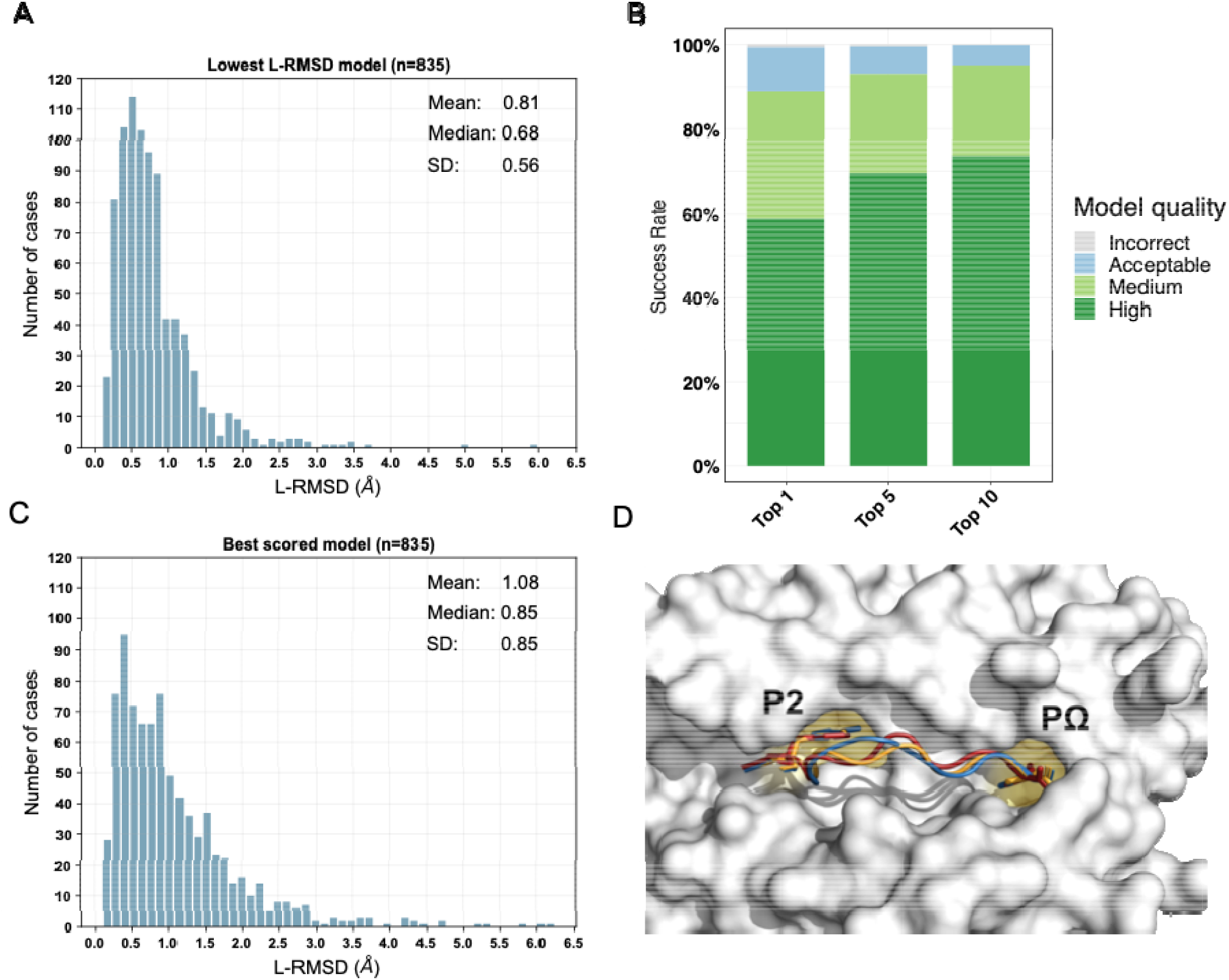
The benchmark performance of PANDORA on 835 pMHC-I complexes with X-ray structures. **A)** Sampling performance of PANDORA. Histogram of the lowest backbone L-RMSD models is shown. **B)** Success rate of Backbone L-RMSD at different thresholds according to CAPRI criteria: High-quality (L-RMSD <1 Å), Medium, (<2 Å), Acceptable (<5 Å), and Incorrect (<10 Å) (Lensink et al., 2020). **C)** Complete performance of PANDORA (modelling + scoring). Histogram of the backbone L-RMSD of the best molpdf models is shown. **D)** Example of an average-quality 3D model generated with PANDORA. The target peptide (PDB ID: 3I6L) is marked in red; the template structure (PDB ID: 3WL9) is marked in blue; the model structure is marked in orange.

To obtain an estimation of which results a user can expect given information known a priori, we checked the performance of PANDORA with respect to different peptide lengths (Suppl. Fig. 2A,B), sequence identities between query and template peptides (Suppl. Fig. 2C,D) and MHC allele difference between target and template (Suppl. Fig 2E,F). PANDORA gives the best performance on 8-9 mer peptides with an average L-RMSD of 0.67 L. PANDORA models generated with 100% peptide identity are slightly better than other peptide similarities (0.61 Å in median RMSD), but no clear trend is observed with respect to the peptide sequence identity (Suppl. Fig. 2C,D). Reasonable performances were also reached in the rare cases (16 out of 835) in which no template from the same gene as the target was available (Suppl. Fig. 2E,F). This shows how PANDORA can be used to build model cases of well-known as well as rare alleles.

### 2.3 PANDORA performs competitively with state-of-the-art methods

We compared PANDORA with three existing methods for pMHC-I 3D modelling: DockTope (30), GradDock (29) and APE-Gen (27) on datasets used in their publications. As not all methods used scoring functions to select best models for their experiments, we compared with each method’s best scenario. Specifically, we used our top molpdf model (PANDORA’s default user output) to compare with the pipelines that used scoring functions (GradDock and DockTope), and our models with best L-RMSDs to compare with the pipeline that reported the best L-RMSD models (APE-Gen).

As shown in Figure 3, PANDORA is competitive with the state-of-the-art methods in terms of best-generated and top-selected models, both lower L-RMSD on average than the ones produced by the published methods. The comparison between PANDORA and GradDock (Figure 3A) and DockTope (Figure 3B) are based on cross-docking. The comparison in Figure 3C reflects the results from the cross-docking experiments from PANDORA and the self-docking experiments from APE-GEN. Whereas self-docking uses the original bound conformations of target MHC and peptide as input to their modelling protocol, cross-docking inputs instead consist of conformations of MHC and peptide that are not the target one. Therefore, self-docking experiments are a simpler scenario than cross-docking experiments, and tend to give better results (29).

**Figure 3.**
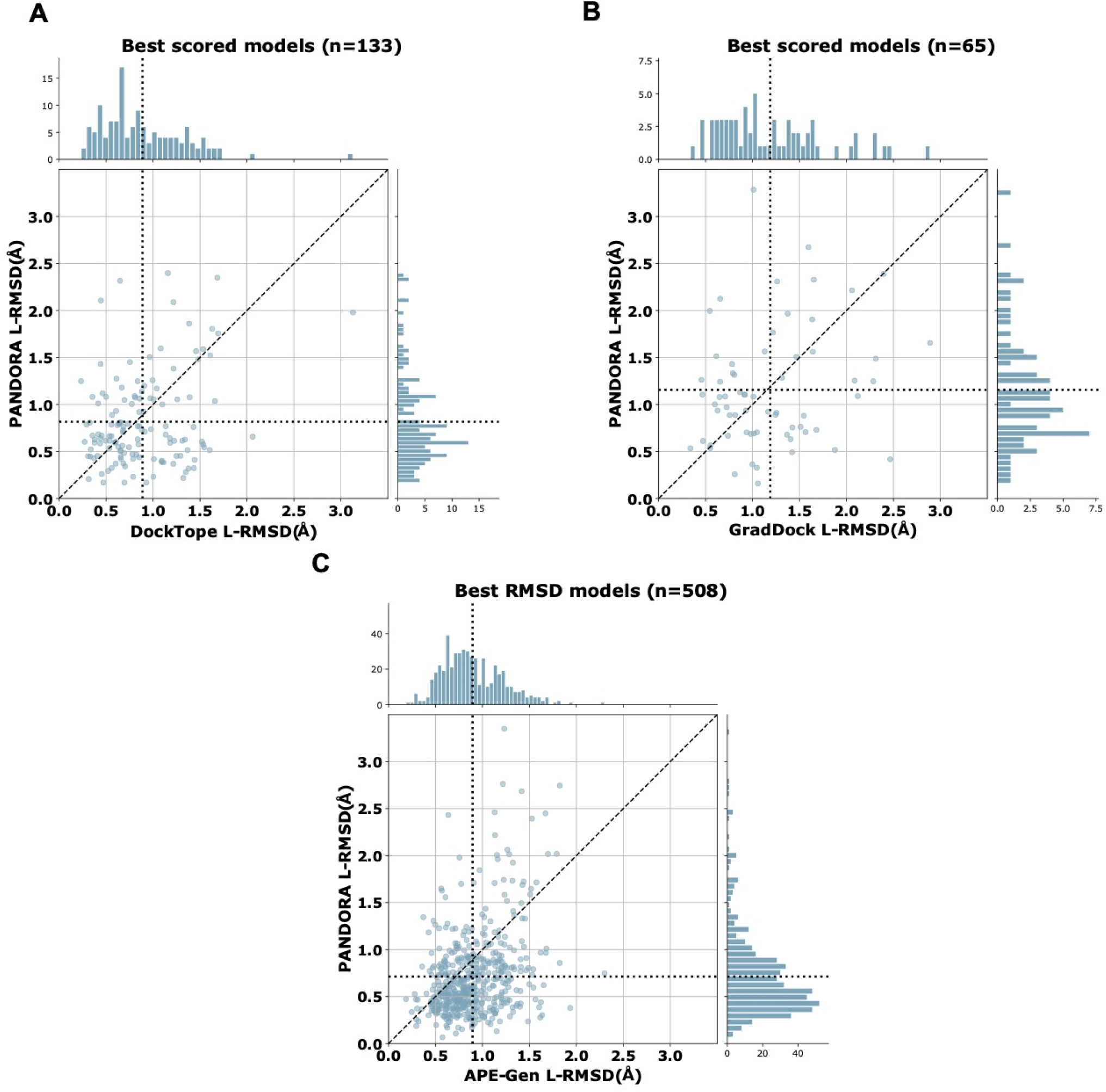
PANDORA comparison with the state-of-the-art methods. Y axes represent PANDORA L-RMSD per case while X axes represent the L-RMSD of the same case modelled with the reported method. The dotted line indicates the average L-RMSD for each method. A) Difference between PANDORA best molpdf model ClIJ L-RMSD and DockTope reported ClIJ L-RMSD on 133 cases (PANDORA cross-docking against DockTope cross-docking); B) Difference between PANDORA best molpdf model backbone + C/J L-RMSD and GradDock reported backbone + C/J L-RMSD on 65 cases (PANDORA cross-docking against GradDock cross-docking); C) Difference between PANDORA best L-RMSD model ClIJ L-RMSD and APE-Gen reported ClIJ L-RMSD on 509 cases (PANDORA cross-docking against APE-Gen self-docking).

PANDORA is computationally efficient. After downloading or building the templates dataset (both options have to be done only once, but building can require up to 1.5 hours) PANDORA takes an average of ~2.6 minutes (156 seconds) to build 20 models per each case on one thread on a Intel(R) Xeon(R) Gold 6142 CPU @ 2.60GHz. According to their publications, DockTope takes “less than 6 hours”, and GradDock takes about “107.79” seconds to model one case, but lacking the exact hardware information a fair comparison is not possible. Given the availability and installation conditions of the softwares we compared with, a direct comparison of the running times can in fact be done only with APE-Gen. APE-Gen takes 3 minutes to prepare the MHC 3D structure plus 2 minutes per each pMHC-I complex using 6 or 8 threads (41). With roughly the same computational time and number of cores (i.e., 5 minutes on 6 to 8 cores), PANDORA can model up to 11-15 cases.

To have a qualitative evaluation of PANDORA against AlphaFold, we tested multiple published strategies for protein-peptide interaction modelling. We tried multimer-approach (36) and linker-approach (37,38) using template-based and template-independent AlphaFold publicly available colabs (34,42). As reported in Suppl. Fig. 3, PANDORA always generated models with a considerably lower backbone L-RMSD than AlphaFold on the four randomly selected pMHC complexes. Also, PANDORA’s cost in terms of computational resources previously discussed is much lower than that of AlphaFold, which can take up to 18 GBs of GPU power for 20 minutes to model one single pMHC case, making modelling of millions of pMHC very expensive with such a tool.

### 2.4 Correct anchor positions play a key role

As mentioned in section 2.1, the input we provided PANDORA with was the actual target peptide anchors that were calculated directly from the target structure. We did so to avoid biases derived from wrong anchor prediction in our benchmark performance. The anchor information is crucial for our modelling pipeline. The majority of 9- to 12-mer MHC-I peptides have canonical anchoring positions at P2 and PΩ (10,11). To study the effect of non-canonical peptide anchoring in the 3D modelling process of PANDORA, we listed which peptides from our benchmark dataset used non-canonical anchor positions to bind to the MHC, resulting in a total of 34 cases. We modelled them as in the previous benchmark experiment, with the only difference that canonical anchor positions were used as input of PANDORA. The models with the lowest L-RMSD are reported in Figure 4A, where the average L-RMSD improvement achieved using real anchors over canonical anchors is 1.48 L. This result is better exemplified in Figure 4B, where it can be observed how defining incorrect (canonical, in this case) anchors causes PANDORA to fix the wrong residues inside the anchor pockets, stretching (as shown) or elongating the peptide central loop, thus worsening the L-RMSD with the x-ray structure.

**Figure 4.**
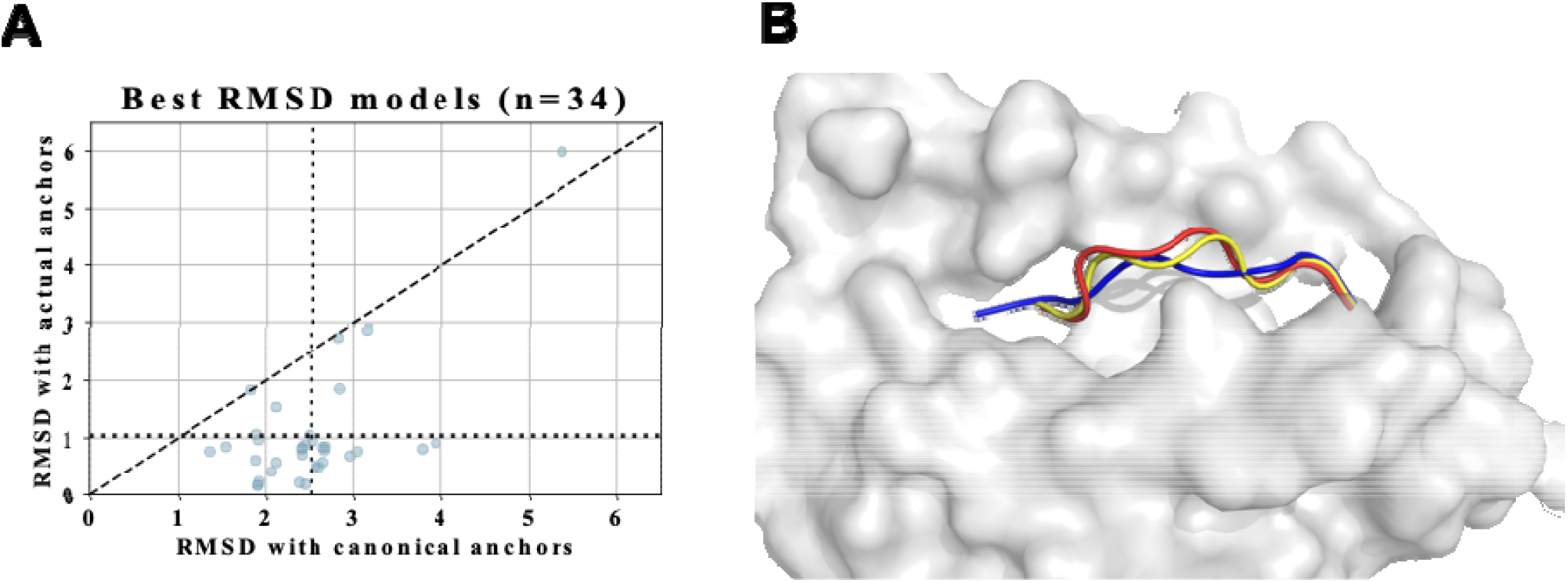
PANDORA’s performance on 34 cases with non-canonical anchor positions. A) PANDORA produced better models than using canonical anchor positions in terms of backbone L-RMSD of cases. B) A typical case (target PDB ID: 1DUY, template PDB ID: 1AO7, peptide=LFGYPVYV) with non-canonical anchor positions (actual anchors: P1 and P9(Ω), canonical anchors: P2 and P9). Red: target peptide. Orange: model with actual anchors. Blue: model with canonical anchors.

Since the real anchor position is hardly available with peptide-MHC binding data, we evaluated the reliability of predicting anchor positions using prediction tools. NetMHCpan4.1 is a binding affinity and core prediction tool for pMHC-I complexes (17). An overview of NetMHCpan4.1 performance on predicting the right anchors over our whole dataset can be found in Suppl. Fig 4, showing that NetMHCpan4.1 provides reliable anchor predictions for most of the cases: correct anchor prediction in 96.5% of the tested cases, one residue shift for 3.2% cases, and position shifts on both anchors in 0.2% cases. Based on these observations, we implemented the following options in PANDORA: to use either canonical anchors, NetMHCpan4.1 predicted anchors, or anchor points provided by the user, who may exploit different tools or gather integrative information from experimental data.

### 2.5 Long peptides are challenging to be modelled reliably

Long peptides (11-15 mers) present a challenge to be modelled reliably (Suppl. Fig. 2A). This is because long peptides are able to fold into small elements of secondary structure in their central part. This problem, although rare (17 cases out of 835 structures in our benchmark dataset presented elements of secondary structure), presents a modelling challenge (model L-RMSD up to 3.03 Å).

To address this challenge, we enabled PANDORA to include secondary structure restraints and tested its performance on a 10-mer (PDB ID: 3BEW) and a 15-mer case (PDB ID: 4U6Y). In these structures the peptide presents in its center region a small alpha helix (16 structures out of 835) and a small beta-sheet hairpin (1 structure out of 835), respectively. We manually defined secondary structure restraints for the peptide based on the bound conformation found in the PDB struct re.

Secondary-structure restraints improved model qualities for both cases (Figure 5). This clearly shows that a correct secondary structure prediction can effectively be provided to PANDORA to guide its modelling, leading to high-quality models even for challenging cases.

**Figure 5.**
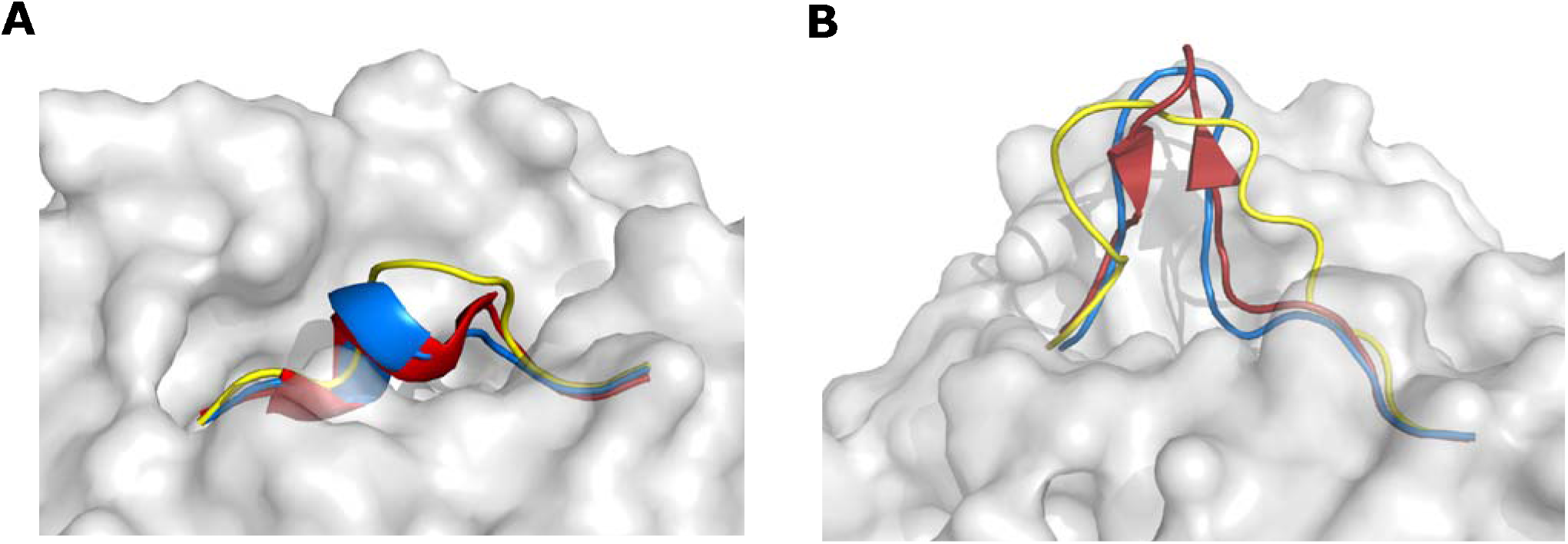
Case study on two long peptides with elements of secondary structure. Red: target peptide. Blue: peptide modelled with secondary structure restraints. Yellow: peptide modelled without secondary structure restraints. The images are oriented to present the most representative view of the difference between models and target. **A)** Case study on the 10-mer from PDB structure 3BEW. L-RMSD with default settings: 2.02 D; L-RMSD using secondary structure restraints: 0.80 D. **B)** Case study on the 15-mer from PDB structure 4U6Y. L-RMSD with default settings: 3.32 D; L-RMSD using secondary structure restraints: 1.50 D.

### 2.6 Software Information

PANDORA is designed to be a robust and user-friendly python package, which can be integrated into other python pipelines. It is highly modularized (see Suppl. Fig. 5 for the object relation diagram) and supports continuous integration, facilitating automatic integration of code development from multiple developers.

PANDORA builds its template database through a robust, automated and yet adjustable module. This module takes care of downloading, parsing and homogenizing the queried types of structures and it summarizes their information (e.g., sequences, allele information, anchor positions, biopython structure object) in an easily accessible python object, providing the base for other methods that might use these data for different purposes. The same module also downloads and parses reference sequences from the manually curated sequence database https://www.ebi.ac.uk/ipd/mhc/ (43,44) to build a local, reliable MHC-I sequence database of multiple species. Both the structural and the sequence database can be rebuilt or updated at any moment with ease by the user, and multiple databases with different parsing criteria can be saved at the same time.

PANDORA takes as input: 1) peptide sequence and 2) MHC allele name. PANDORA by default assumes the canonical anchor positions. Users may easily personalize their runs by adding anchor information or secondary structure predictions, increasing the number of generated loops, changing refinement mode or providing personalized MHC sequences. Expert users of MODELLER may further personalize the main MODELLER modelling scripts to adapt the pipeline to their specific needs.

PANDORA is computationally efficient (see Section 2.1 for average running times) and copes well with large-scale modelling tasks on HPC (High Performance Computing) facilities. PANDORA supports parallelization at multiple levels (per-case or per model). A short tutorial with six different examples showing the ease of setting up different types of PANDORA run can be found in our GitHub repository (https://github.com/X-lab-3D/PANDORA). Users can report problems, ask for assistance or request specific features to be added from the GitHub issue section.

## 3 Discussion

In this study we present PANDORA, a reliable pMHC modelling pipeline. To reduce the computational time associated with the modelling, PANDORA exploits two pieces of domain knowledge: 1) MHC structures are highly conserved, 2) MHC uses anchor positions to bind the peptide.

We demonstrate here that PANDORA performs reliably on the largest pMHC-I dataset obtained from IMGT/3Dstructure-DB (39) and used for cross-validation. PANDORA also shows competitive performance compared with three state-of-the-art modelling softwares while PANDORA outperforms them in computational efficiency and ease of installation. Being a modular python package, our method (or single sub-modules of it) can be integrated in other pipelines with ease, which is considerably harder for the state-of-the-art softwares.

Although specific software for pMHC modelling are available, we could not avoid to compare our software with the groundbreaking, general-purpose 3D-modelling software AlphaFold (34), that has recently been used to accurately model tens of thousands of protein structures (45) including MHCs. PANDORA outerforms AlphaFold on the pMHC modelling task in terms of accuracy and computational time. Our evaluation of AlphaFold on modelling pMHC-I complexes revealed that AlphaFold is often misplacing the P2 anchor residue outside its pocket, causing a high backbone L-RMSD compared to the X-ray structure (Suppl. Fig. 3A,B and C). Furthermore, in presence of secondary structures, AlphaFold can generate even higher L-RMSD models (Suppl. Fig. 3D).

When run with default settings (i.e., using P1 and PΩ as anchors for pMHC-I), our method achieves a median L-RMSD of 0.67 Å on our large benchmark dataset (Suppl. Fig. 6), but it fails in delivering high-quality models for some, mainly non-canonical cases. Most of these outliers are caused by peptides with non-canonical anchor positions. To overcome this issue, users may opt for binding core prediction tools (such as NetMHCPan4.1 (17) or MHCflurry (16)) to guide PANDORA’s prediction or model peptides with multiple anchor positions and choose the ones with best molpdf scores. Also, high L-RMSD can be caused by long peptides able to fold into secondary structures. In these cases, users may decide on secondary structure- or folding-prediction tools such as AGADIR (46) and PEPFOLD3 (47) to elicit secondary structures restraints to input into PANDORA. For long peptides that do not form secondary structures, users might just use a much larger sampling step, increasing the number of generated models to hundreds or thousands. We report in Suppl. Fig 7 how a larger sampling results in slightly increased models quality.

While we evaluated PANDORA on MHC-I cases only here, PANDORA can be directly applied to MHC-II. MHC-II mediated T cell responses account for the predominant response upon cancer vaccinations (3,48,49). We systematically investigated PANDORA’s performance on pMHC-II (manuscript under preparation). PANDORA with support for both MHC-I and MHC-II is freely accessible for academic usages (see Code Availability).

PANDORA supports multi-level parallelization and multiple user-configurable options (see github README at: https://github.com/X-lab-3D/PANDORA). These features make it suitable for high-throughput purposes as well as to explore the modelling of particularly challenging peptides (e.g., peptides of non-canonical length). In fact, its computational efficiency allows users to quickly run thousands of cases or to increase the models’ sampling (from the default of 20 to hundreds or thousands) to explore a higher variety of conformations. This computational efficiency is combined with easy installation, flexibility, robust template data collection and high quality of the produced models. PANDORA thus makes a reliable tool for research groups that might need either fine-tuned, accurate 3D models of single pMHC cases, or large-scale modellings (both on HPC facilities or modest desktops).

Lastly, PANDORA is able to enrich the large amount of existing sequence-based binding data with high-quality 3D models, providing 3D enriched data to subsequent ML algorithms. Considering PANDORA’s accuracy and computational requirements in fact, it can be used to affordably generate millions of 3D models. 3D-based AI frameworks like DeepRank (51) and DeepRank-GNN (52) can then exploit these to tackle long standing challenges in pMHC-based vaccine design (work in progress).

## 4 Materials and Methods

### 4.1 PANDORA protocol

The PANDORA package generates pMHC 3D models through restraint-guided homology modelling. PANDORA can take as input one or multiple peptide sequences and an MHC-I IMGT allele name for each peptide. It returns by default 20 model structures (adjustable) in PDB format, ranked by MODELLER’s internal scoring function molpdf (32). To build the pMHC models, PANDORA covers three main steps (shown in Figure 1C): i) template set building, ii) input preparation and iii) 3D modelling, described below. Although PANDORA works for MHC class I and II, below we focus on the protocol for pMHC-I as our experiments presented in this paper are on MHC-I (MHC-II manuscript under preparation).

#### 1) Template Set Building

PANDORA automatically builds an extensive cleaned template set. Template pMHC-I structures with peptide length spanning from 7 to 15 (adjustable) residues are downloaded from the IMGT/3D-structureDB (39). From each PDB file only one Alpha chain (if multiple copies are available) and its bound peptide are extracted (β2-microglobuline is neither saved in the template object, nor modelled). If present, non-canonical residues are changed into canonical residues when no coordinate modifications are required (e.g., changing phosphoserines in serines by removing the phospho group) (see Suppl. Table. 2 for the list of tolerated non-canonical residues). However, the template is removed from the dataset when: 1) other non-canonical residues are present; 2) a small, non-amino acid molecule besides the peptide is present inside the MHC binding cleft; 3) the PDB structure cannot be parsed in Biopython (53) for additional reasons; or 4) the file is lacking allele information from IMGT. The parsing resulted, for our cross-validation benchmark experiment, in a total of 1188 pMHC-I PDB downloaded structures (as of March 23rd of 2021) and a final parsed dataset of 835 PDB structures over 78 MHC I G-domain alleles (the case 3RGV had to be manually removed due to unexpected issues in the parsing).

#### 2) Input Preparation

##### Template selection

For each pair of MHC allele type and peptide sequence provided by the user, PANDORA searches the template database, computes a list of putative templates and selects the first of the list as template. First, it searches for templates that share the same MHC allele type as the target. If no such templates are available, the list is compiled with templates from the same allele group. If these do not yield any templates either, PANDORA extends the search for templates from the same gene. Putative template peptides are then aligned with the target peptide by making sure the anchors are aligned and gaps (if needed) are added in the center of the flexible loop. These alignments are ranked with a PAM30 matrix to select the best template.

##### Alignment File generation

Once a template is selected, its MHC sequence is aligned with the target by using MUSCLE (54), while for the peptides the dummy alignment generated for the template selection step is maintained. For the benchmark experiment, the MHC sequence for each case was retrieved from the target structure to be modelled. Besides MHC types, users may also provide MHC sequences. If a user does not provide MHC sequences, PANDORA will automatically retrieve it from the reference MHC allele sequence (retrieved from https://www.ebi.ac.uk/ipd/mhc/ (43)) according to the provided allele name.

#### 3) 3D Modelling

##### MHC Homology Modelling

PANDORA is built on top of MODELLER (32). The template structure file, anchor positions and the template-target alignment file are fed into MODELLER to generate target pMHC models. First, the MHC structure is generated with a simple homology model over the template structure. Then, anchor positions are provided to MODELLER to indicate which part of the peptide should be kept restrained. Finally, the central part of the peptide is refined with a short energy minimization. Generated models are then ranked using MODELLER’s built-in molpdf function for selection of near-native decoys. In case the target peptide sequence and MHC allele are identical between target and template, the initial loop model generated by MODELLER (which has the same structure as the template) is scored as top model (with a dummy molpdf score) and provided as best output. Also, the user is informed of this sequence identity and pointed to the deposited X-ray structure from PANDORA’s log.

### 4.2 Comparisons with state of the art

Datasets for comparisons with state-of-the-art methods were retrieved from each software’s paper or kindly provided by the authors. Some structures could not be processed by PANDORA according to the criteria listed in section 5.1 or were not found in IMGT/3Dstructure-DB, resulting in smaller comparison datasets than the exact ones provided in literature. We used 133 out of 135 structures for Figure 3A (DockTope), with a peptide length span from 8 to 10 over 5 MHC alleles; 65 out of 69 for Figure 3B (GradDock), with a peptide length span from 8 to 10 over 21 MHC alleles; and 508 out of 535 for Figure 3C (APE-Gen), with a peptide length span from 8 to 11 over 59 MHC alleles.

### 4.3 Evaluations

#### Ligand Root Mean Squared Deviation (L-RMSD)

The models’ quality was evaluated in terms of L-RMSD (40). To this purpose, the G-domains (positions 1-180) of models and target structures were superposed and the L-RMSD was calculated as the RMSD between the atoms of the experimentally determined peptide conformation and the modelled peptide. L-RMSDs were calculated using profit (55). To directly compare with state-of-the-art methods, we used the same sets of atom as these works did: Carbon α L-RMSD (for DockTope and APE-Gen, Figures 3A and 3C respectively) and Backbone + Carbon fJ L-RMSD (for GradDock, Figure 3B).

#### Hit Rate and Success Rate

Hit Rate and Success Rate are widely used in computational modelling for biomolecular complexes (56). A hit here is a model with an L-RMSD < 2 Å from the target structure (57). The Hit Rate is defined as the percentage of hits taken when selecting the top N ranked models, averaged over every case:

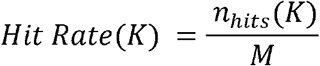

where *n*_*hits*_(*K*) is the number of hits (i.e., near-native models) among top K models and *M* the total number of near-native models for this case.

The Success Rate is defined as the number of cases, taken the top N ranked models, containing at least one hit, divided by the total number of cases:

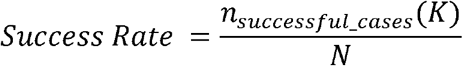

where *n*_*successfull_cases*_(*K*) is the number of cases with at least one hit among top K models, and N is the total number of cases.

## 5 Data & code availability

The list of PDB structure IDs used in the cross-validation benchmark experiment can be found in Supplementary Table 1. The PANDORA database used for the experiments in this work is available for download at: https://github.com/X-lab-3D/PANDORA_database. Related data will be deposited to SBGrid with DOI upon acceptance of the manuscript.

The source code is available on GitHub: https://github.com/X-lab-3D/PANDORA and the package will be released to PyPI (the Python Package Index) at https://pypi.org/project/csb-pandora/ and Zenodo upon acceptance of the manuscript. PANDORA is implemented in Python 3.7 and leverages on Biopython (53) and MODELLER (32). PANDORA and all the required packages can be easily installed as shown by our GitHub README file (https://github.com/X-lab-3D/PANDORA/blob/master/README.md), but the user will still be requested to input a MODELLER license code (freely available for academic purposes at https://salilab.org/modeller/registration.html).

## 6 Conflict of Interest

The authors declare that the research was conducted in the absence of any commercial or financial relationships that could be construed as a potential conflict of interest.

## 7 Author Contributions

LX, DFM and FMP contributed to the design of the pipeline and the design and development of the experiments. FMP and DS contributed to the preliminary experiments. DFM, DVT, FMP, RB and NR contributed to the development of the pipeline. NR, DFM and FMP contributed to the release of the python package and online documentation. DFM, FMP, LX and DVT contributed to the writing of the manuscript.

## 8 Funding

This project is supported by the Hypatia Fellowship from Radboudumc (Rv819.52706).

## 9 Acknowledgments

The authors thank Prof. Hak-Sung Kim, Dr. Yoonjoo Choi and Dr. Hyun-Ho Kyeong for kindly providing data and information about GradDock, Prof. Lydia Kavraki for kindly providing the data for comparison with APE-Gen, Dr. Maurício Menegatti Rigo for kindly providing the data for comparison with DockTope. We also thank Prof. Alexandre Bonvin and Prof. Peter-Bram ‘t Hoen for experimental and writing advice and Dr. Siri C. van Keulen for writing advice.

## Notes

https://github.com/X-lab-3D/PANDORA

https://github.com/X-lab-3D/PANDORA_database

### Competing Interest Statement

The authors have declared no competing interest.

