## Supplementary Materials for "PANDORA: a fast, anchor-restrained modelling protocol for peptide:MHC complexes"

### Supplementary Figures

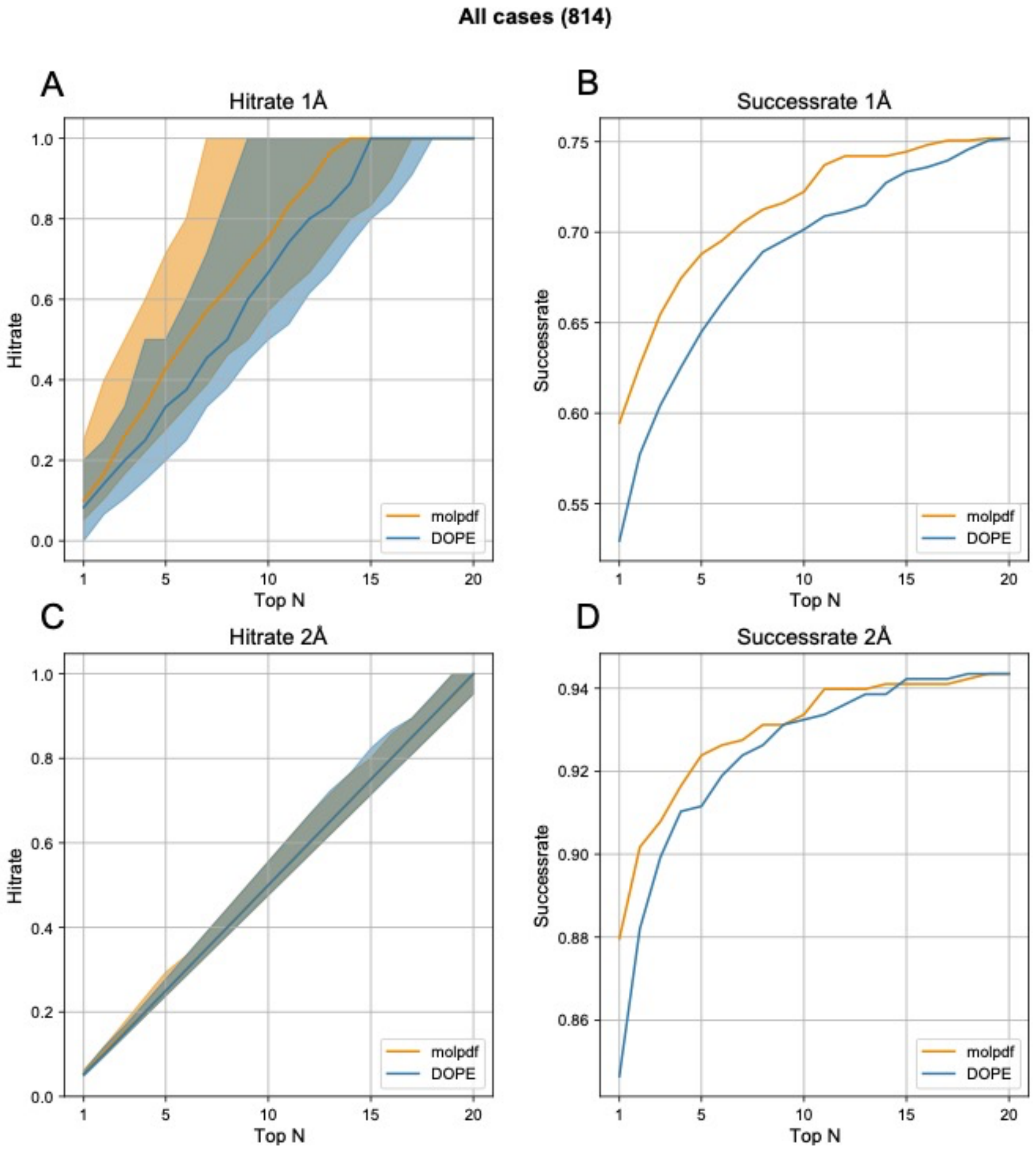


**Supplementary Figure 1. Comparison between MODELLER's internal scoring functions molpdf and DOPE.** For the Hit rate plots, the line marks the average of the Hit Rates (see Methods) and the shaded area marks the 25%- 75% quantile interval. For the "1Å" plots (A and B) a hit is defined as a model with backbone L-RMSD <= 1 Å, while for the "2Å" plots (C and D) a hit is defined as a model with backbone L-RMSD <= 2 Å . Note that these results were obtained on an earlier experiment on of 814 cases. **A)** Hit rate at 1Å **B)** Success rate at 1 Å **C)** Hit rate at 2 Å **D)** Success rate at 2 Å.

**All cases (835)**

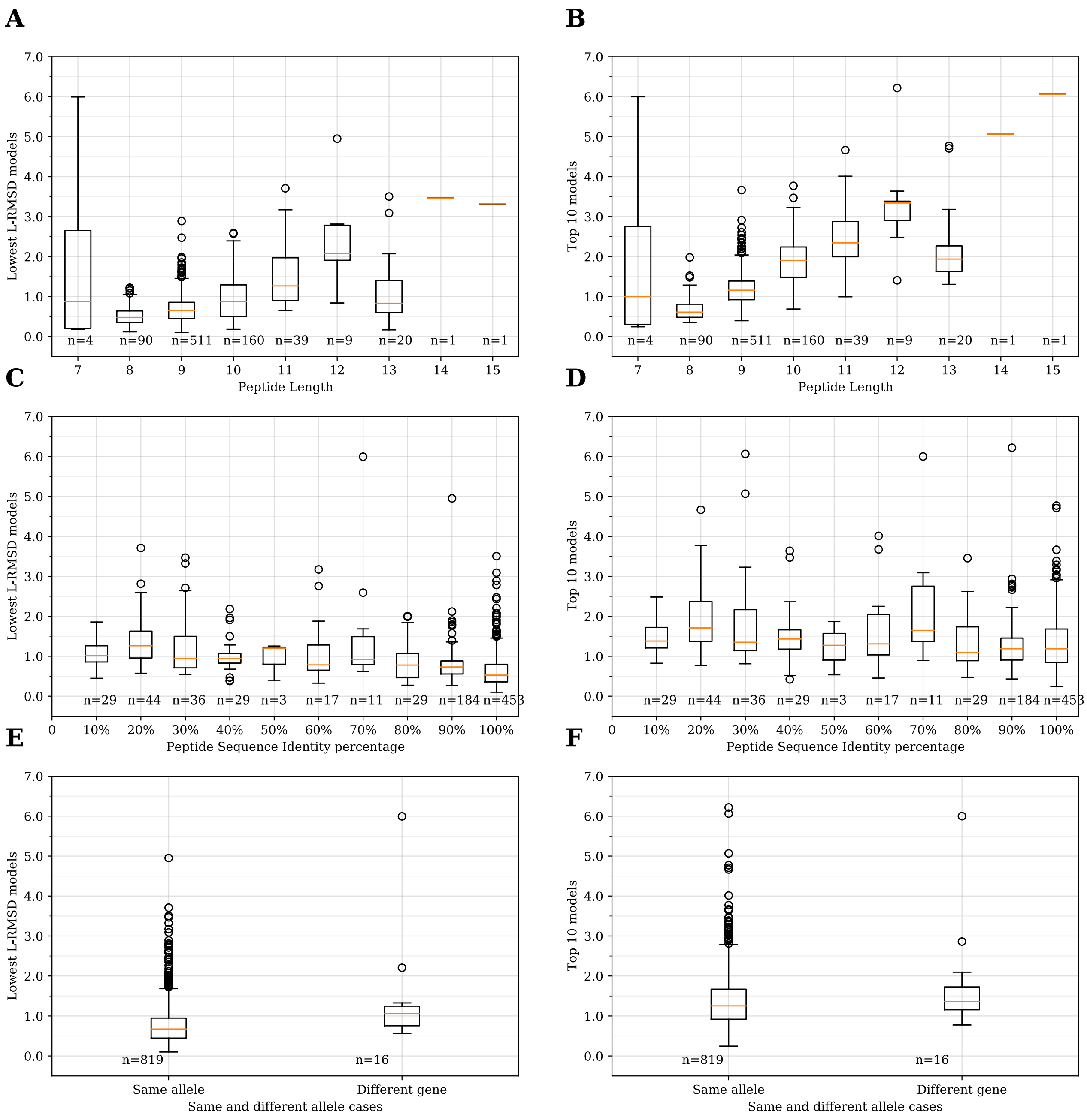


**Supplementary Figure 2. PANDORA benchmark results with respect to peptide length, peptide sequence identity and MHC allele types.** Left panels (A, C and E) represent the best models generated by PANDORA among the top 20 models (without the scoring step). Right panels (B, D and F) represent the top 10 models (with the scoring step). **A and B)** PANDORA's performance over peptide length. **C and D)** PANDORA's performance over target-template peptide sequence identity. **D and F)** Using templates with different MHC types vs. with the same MHC types as the target.

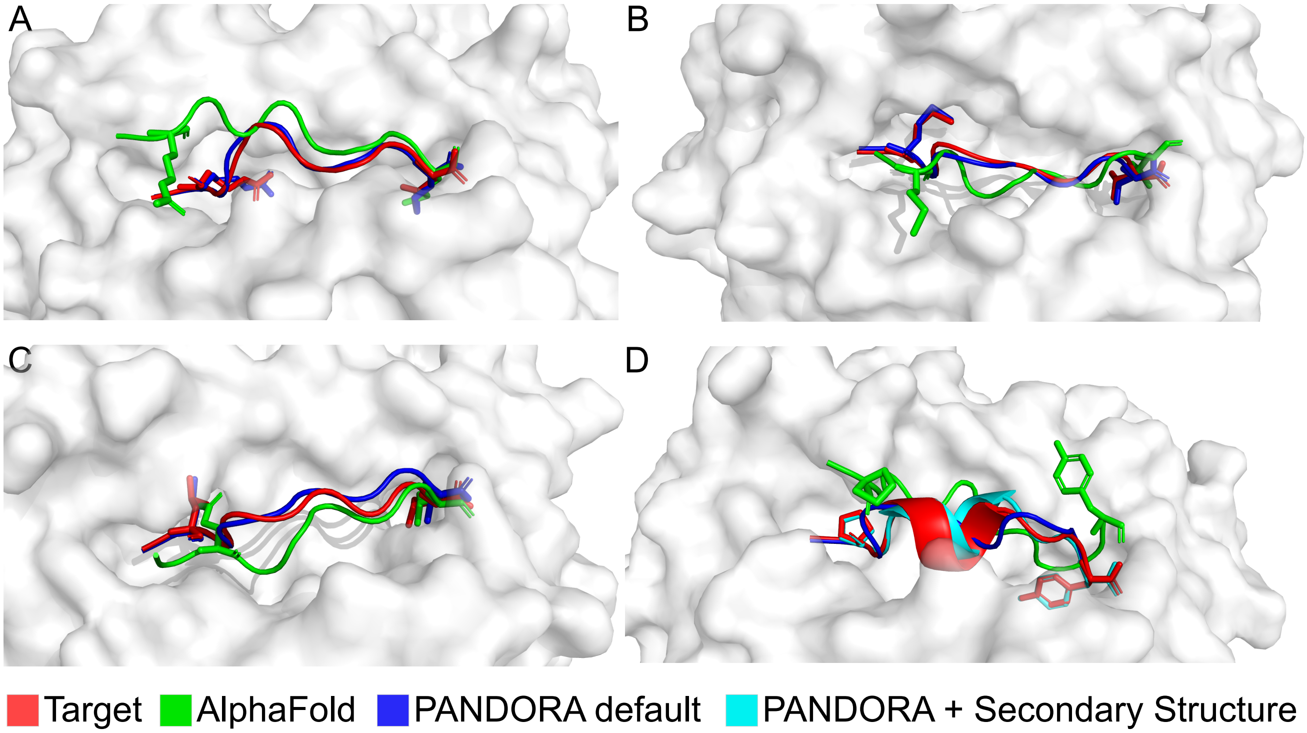


**Supplementary Figure 3. Model quality comparison of AlphaFold and PANDORA**. Red: target peptide; Green: peptide modelled with AlphaFold (best model); Blue: peptide modelled with PANDORA (best of top 5 molpdf); Cyan: peptide modelled with PANDORA adding secondary structure restraints. The images are oriented to present the most representative view of the difference between models and target. **A)** AlphaFold model generated using AlphaFold_multimer (template independent). PDB structure: 1G7P. AlphaFold backbone L-RMSD: 2.32 Å; PANDORA backbone L-RMSD: 0.86 Å. B) AlphaFold model generated by linking the peptide using Poly-Glycine linker (30 Glycines) (template-dependent). PDB structure: 3BZE. AlphaFold backbone L-RMSD: 4.43 Å; PANDORA backbone L-RMSD: 0.89 Å. C) AlphaFold model generated by linking the peptide using Poly-Glycine linker (15 Glycines) (template-dependent but the target structure was not present in AlphaFold training set). PDB structure: 7N1A. AlphaFold backbone L-RMSD: 3.60 Å; PANDORA backbone L-RMSD: 0.94 Å. D) AlphaFold model generated using AlphaFold_multimer (template-independent). PDB structure: 4PRE. AlphaFold backbone L-RMSD: 7.04 Å; PANDORA backbone L-RMSD: 1.86 Å; PANDORA + secondary structure restraints backbone L-RMSD: 1.17 Å.

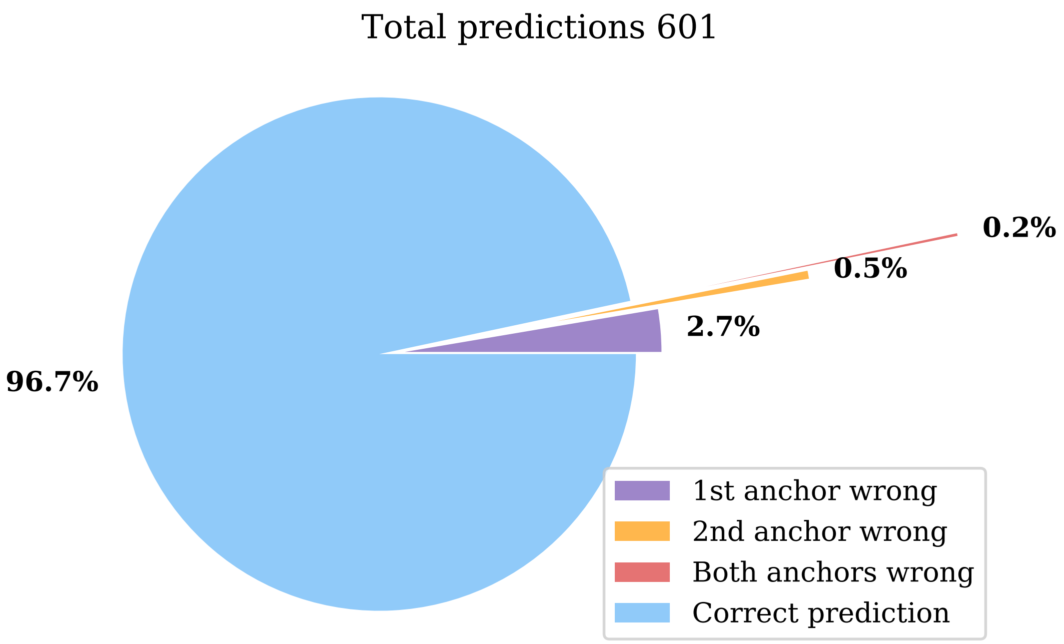


**Supplementary Figure 4. NetMHCpan4.1 anchor prediction performance over PANDORA benchmark dataset.** Upon automatically feeding NetMHCpan4.1 with peptide sequence and allele information from IMGT/3DstructureDB, we were able to obtain 601 predictions out of 835 cases. Missing predictions are mainly due to allele name inconsistencies between NetMHCpan and IMGT sequence databases. Most of the incorrect predictions (17 out of 20) fall between the non-canonical-anchor cases we analyzed in section 2.4. In every case of misprediction, the predicted anchor was only one residue away from the real anchor.

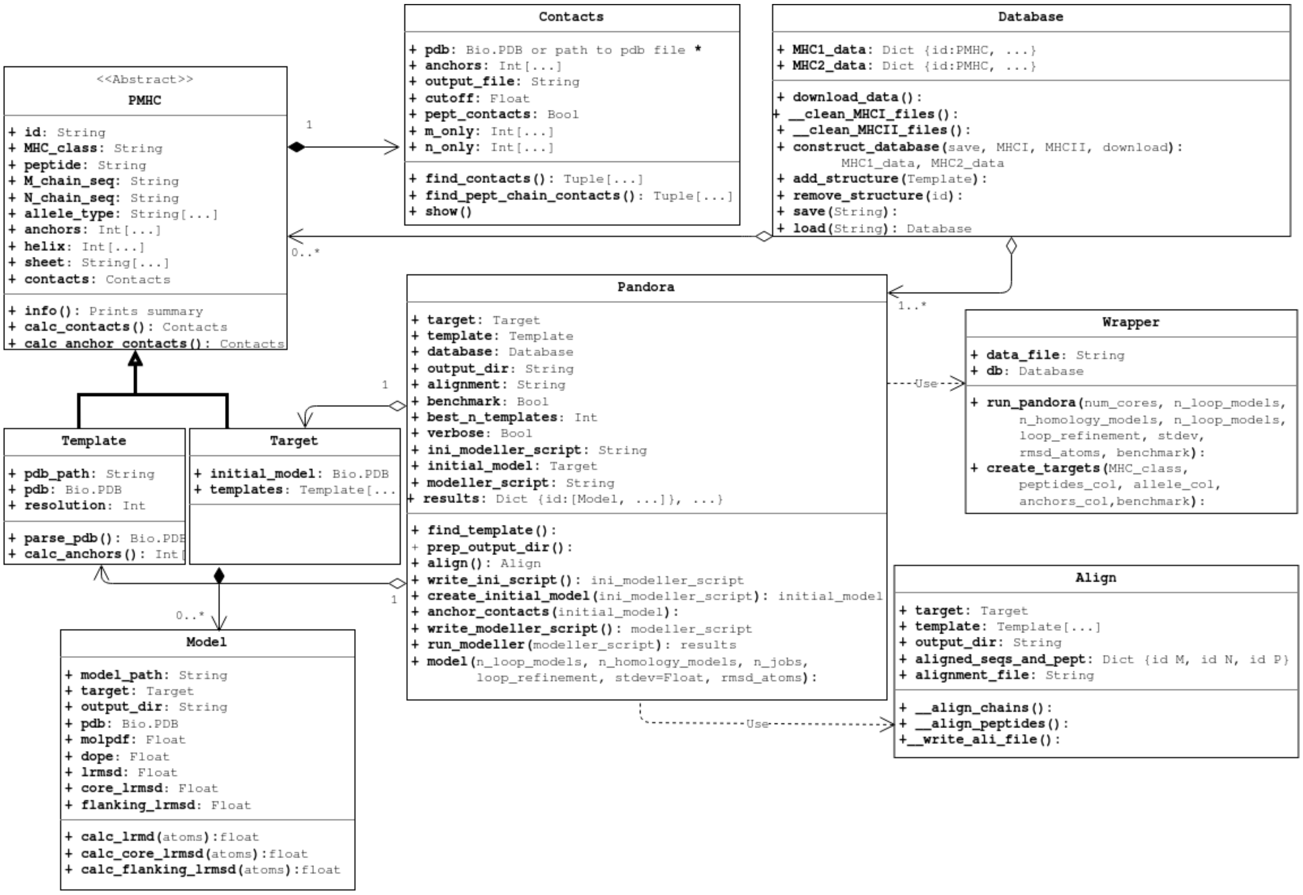


**Supplementary Figure 5. Class diagram for PANDORA's framework.** PANDORA is highly modularized and configurable.

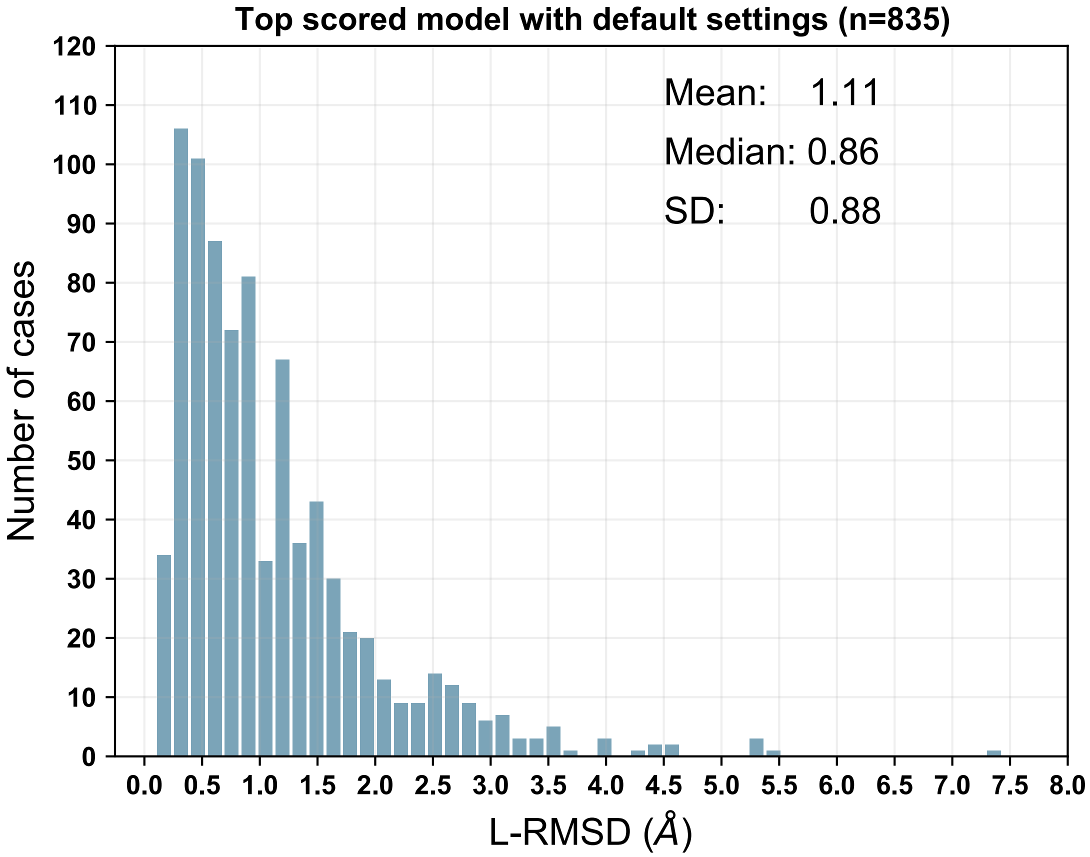


**Supplementary Figure 6. Performance of PANDORA using canonical anchor positions on the benchmark dataset (835 pMHC-I complexes with X-ray structures).** The lowest backbone L-RMSD models are shown.

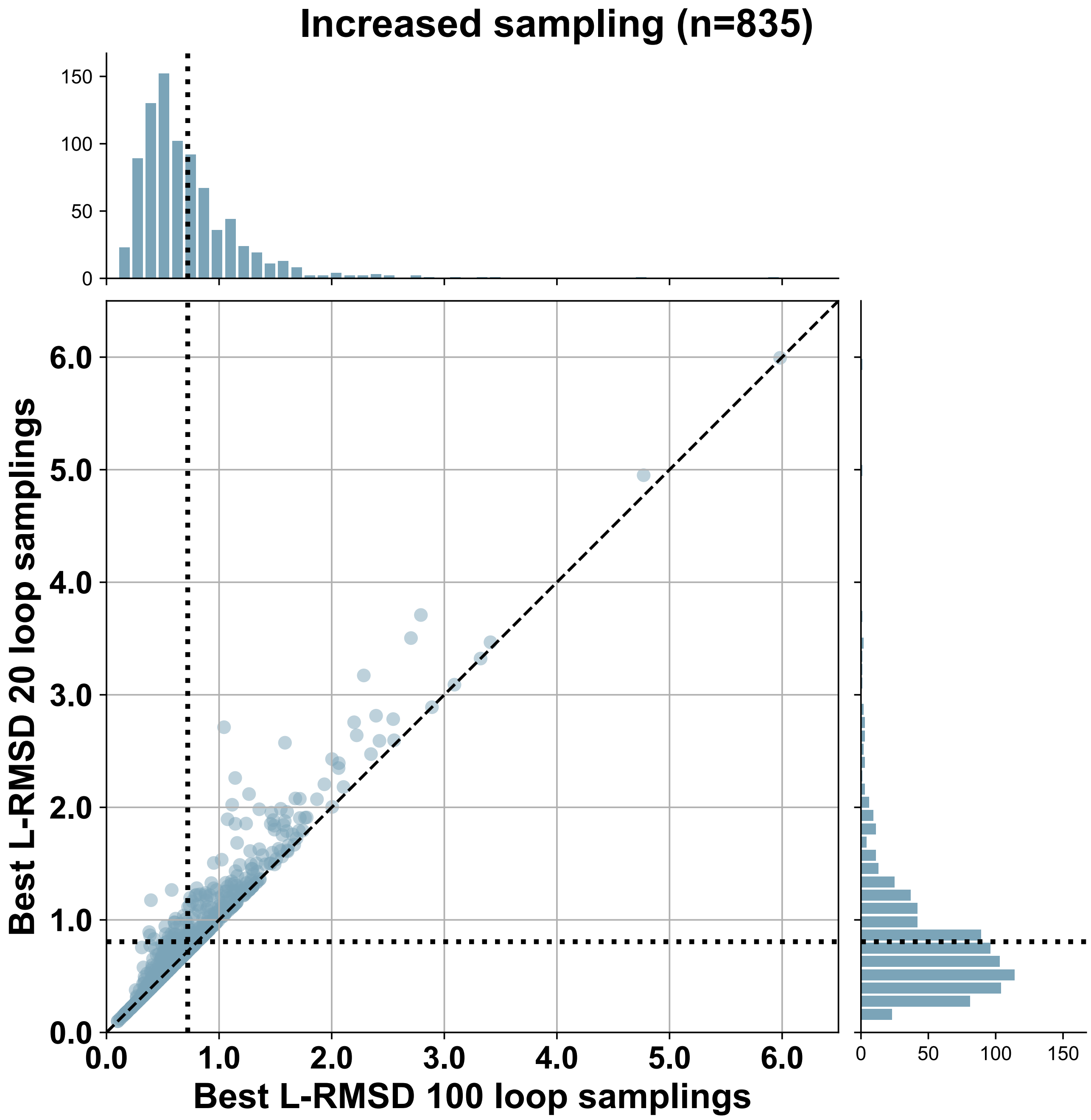


**Supplementary Figure 7. Performance of increased sampling for modelling p-MHCI.** For each case the best backbone L-RMSD is reported. The dotted lines represent the mean of the distribution per axis, the dashed line is the bisector. Actual anchor positions were used.
