## Supplementary Table 2 for "PANDORA: a fast, anchor-restrained modelling protocol for peptide:MHC complexes"

**Supplementary Table 2. List of non-canonical amino acids tolerated by PANDORA parsing step.**

| PDB ID | PDB url link |
|---|---|
| CIR | https://www.rcsb.org/ligand/CIR |
| CSO | https://www.rcsb.org/ligand/CSO |
| F2F | https://www.rcsb.org/ligand/F2F |
| SEP | https://www.rcsb.org/ligand/SEP |
